# Decoupling of evolutionary changes in mRNA and protein levels

**DOI:** 10.1101/2023.04.08.536110

**Authors:** Daohan Jiang, Alexander L. Cope, Jianzhi Zhang, Matt Pennell

## Abstract

Variation in gene expression across lineages is thought to explain much of the observed phenotypic variation and adaptation. The protein is closer to the target of natural selection but gene expression is typically measured as the amount of mRNA. The broad assumption that mRNA levels are good proxies for protein levels has been undermined by a number of studies reporting moderate or weak correlations between the two measures across species. One biological explanation for this discrepancy is that there has been compensatory evolution between the mRNA level and regulation of translation. However, we do not understand the evolutionary conditions necessary for this to occur nor the expected strength of the correlation between mRNA and protein levels. Here we develop a theoretical model for the coevolution of mRNA and protein levels and investigate the dynamics of the model over time. We find that compensatory evolution is widespread when there is stabilizing selection on the protein level, which is true across a variety of regulatory pathways. When the protein level is under directional selection, the mRNA level of a gene and its translation rate of the same gene were negatively correlated across lineages but positively correlated across genes. These findings help explain results from comparative studies of gene expression and potentially enable researchers to disentangle biological and statistical hypotheses for the mismatch between transcriptomic and proteomic studies.

## Introduction

Understanding the causes and consequences of evolutionary divergence in gene expression is important for explaining divergence in organismal phenotypes and adaptation [1]. As proteins carry out functions encoded by protein-coding sequences and are generally thought of as the functional unit of the cell, the protein abundance (hereafter, the protein level) is expected to be the target of natural selection. However, previous work on gene expression evolution has predominantly relied on mRNA levels due to the relative simplicity and cost-effectiveness of high-throughput mRNA sequencing methods compared to mass spectrometry-based proteomics. This implicitly assumes that mRNA levels are an adequate proxy for protein levels; however, many studies have documented weak to moderate correlations (i.e., *<* 0.6) between mRNA and protein levels across genes [2–5], tissues [5–9], and species [10–12]. Disentangling the biological, technical, and statistical explanations for the observed correlations between mRNA and protein levels remains an open and challenging problem [13–16]. In order to understand the biological underpinnings of this relationship — i.e., to better understand how, when, and why discrepancies between mRNA and protein levels arise on evolutionary timescales — we need mathematical models that describe the coevolution between these two aspects of gene expression and that can generate clear predictions.

In this study, we focus specifically on the correlation between mRNA and protein levels of the same gene across species. In addition to a weak to moderate correlation between mRNA and protein levels, protein levels are generally more conserved than mRNA levels across species [10, 11, 17]. This phenomenon is hypothesized to be due to compensatory evolution, in which changes to the mRNA level can be offset by changes to translation regulation, and vice versa [10, 11, 17, 18]. Consistent with the compensatory evolution interpretation is the observed negative correlation between the evolutionary divergence of mRNA levels and translational efficiencies (i.e., per-transcript rate of translation, as measured by ribosome profiling). For instance, the difference between two yeast species, *Saccharomyces cerevisiae* and *S. paradoxus* in the mRNA level and that in translational efficiency of the same gene is more frequently in opposite directions than in the same direction [19, 20]. Similarly in mammals, the amount of divergence across species in the mRNA level and that in translational efficiency are negatively correlated [18]. However, the observed negative correlation between the mRNA level and the translational efficiency may be attributable to a statistical artifact, as translational efficiency estimated using ribosome profiling data is a ratio of the total translation level and the mRNA level, and regressing *Y/X* (or *Y − X* after log transformation) against *X* is well-known to induce spurious negative relationships when there is measurement error in one, or both, of the variables [21–23]. Indeed, findings from other studies find little evidence of compensatory evolution between transcription and translation, with changes to translation largely mirroring changes to transcription [24, 25]. The conflicting observations regarding the coevolution of transcription, translation, and protein levels generally raise the question of the evolutionary conditions likely to result in compensatory evolution and its general prevalence in shaping gene expression evolution.

Complicating biological interpretations from empirical data is a lack of theoretical justification for the model of compensatory evolution between transcription and translation. Compensatory evolution is generally invoked as a post hoc explanation that is based on the assumption that the protein level is the most likely target of natural selection, but it is unknown whether this explanation is adequate to generate the observed patterns under realistic evolutionary scenarios. To address this lack of theoretical justification, we develop a framework rooted in quantitative genetics theory to investigate the coevolution of the mRNA level, the rate of translation, and the protein level of a gene when the protein level is the target of natural selection. Under this framework, we conducted simulations to demonstrate how patterns of evolutionary divergence in these traits and correlation between them are related to values of evolutionary parameters. Our simulations reveal that stabilizing selection on the protein level is sufficient to cause compensatory evolution, which holds across a variety of evolutionary conditions. We also find that evolutionary changes in the mRNA level and that in the rate of translation complement each other when the protein level is under directional selection, resulting in a negative transcription-translation correlation that is similar to that caused by stabilizing selection.

## Results

### Stabilizing selection leads to compensatory evolutionary of transcription and translation

To understand the coevolutionary dynamics of mRNA and protein levels when the latter is subject to natural selection, we considered a model where the mRNA level and the per-transcript rate of translation (more concisely, the “translation rate”) are directly affected by mutations and collectively determine the protein level. The protein level is the fitness-related trait, and we first considered the case when protein levels are subject to stabilizing selection. Under this model, we conducted simulations of evolution in 500 replicate lineages and examined the resulting distribution of phenotypes among these lineages (see Methods). This model assumes mRNA and protein degradation rates remain constant through time, such that all evolutionary changes to mRNA and protein levels are mediated through changes to transcription and/or translation. Here, each replicate lineage can be conceived as a species, and the amount of evolutionary divergence among species was represented by the variance among these lineages. As a negative control, we also simulated mRNA and protein levels when neither trait is subject to natural selection, i.e. neutral evolution.

When the protein level is under stabilizing selection, the correlation between the mRNA level and the translation rate (the “transcription-translation correlation”) were strongly negatively (*r < −*0.95 under the parameter combination we considered, Fig. 1A). In contrast, such a negative transcription-translation correlation was not observed under neutrality (Fig. S1A). The correlation between the mRNA level and the protein level (the “mRNA-protein correlation”) was much weaker under stabilizing selection compared to that under neutrality (Fig. 1B, Fig. S1B). As measurement errors are known to impact empirical measures of gene expression [26], we added noise to each end-point mRNA level and protein level (see Methods) and repeated our analyses to determine if our results were robust. In general, measurement error had little impact on our general results (Table S1). The dependence of measurement error in the translation rate on measurement error in the mRNA level (see Methods) created a trend towards a negative correlation between the mRNA level and the translation rate under neutrality. In contrast, measurement error did not strengthen, but weakened, the negative transcript-translation correlation in the presence of stabilizing selection (Table S1). This is consistent with attenuation bias towards 0, which is known to occur when two correlated traits are measured with error. Variances of both the mRNA level and the translation rate increased over time when the protein level was subject to stabilizing selection, although the variances of each trait were much lower than the neutral expectation (Fig. 1C and Fig. S1C). In contrast, the variance of the protein level saturated early during the simulated evolution in the presence of stabilizing selection (Fig. 1C), which did not happen under neutrality (Fig. S1A).

**Figure 1:**
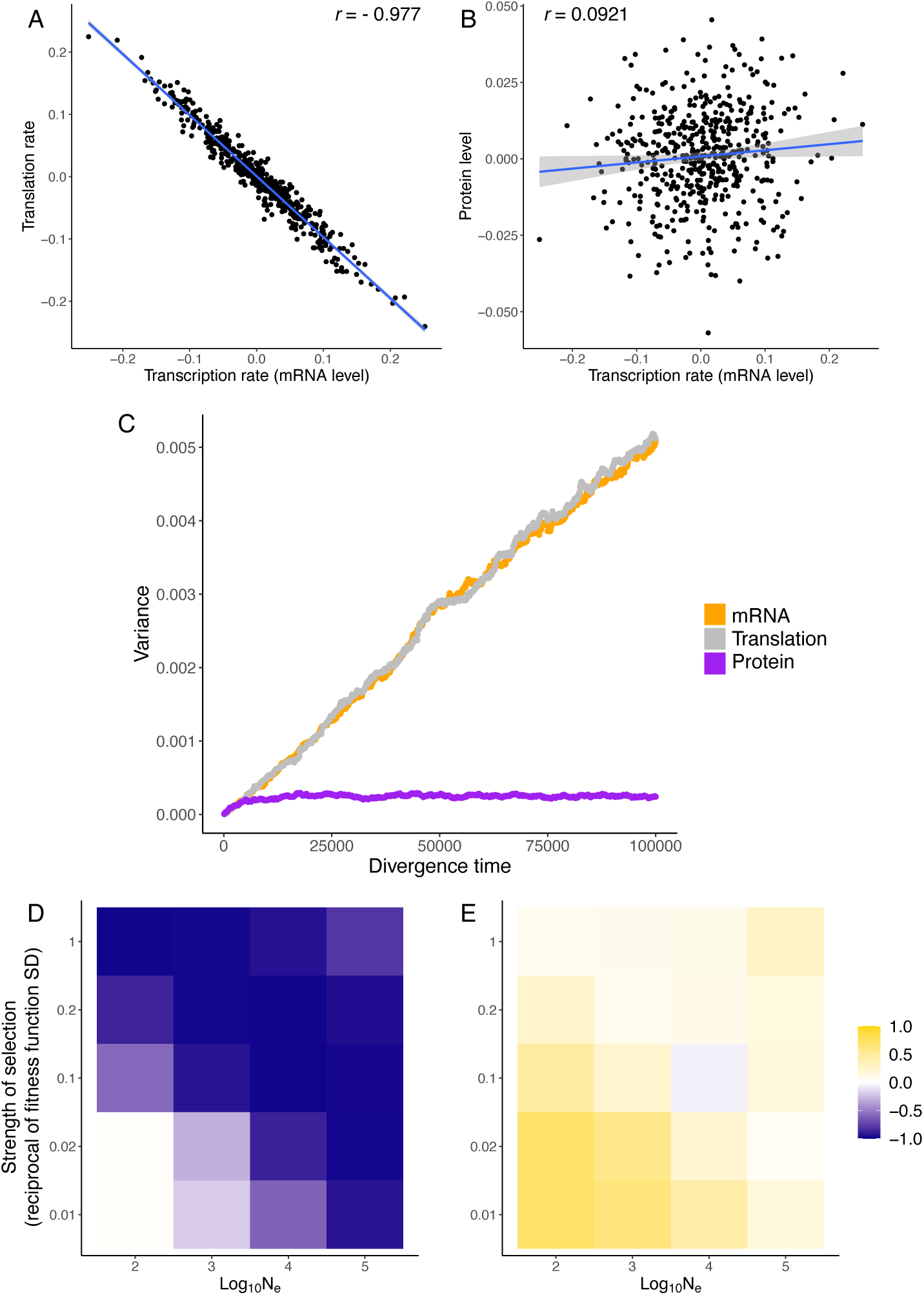
Coevolution of the mRNA level, the rate of translation, and the protein level when the protein level is under stabilizing selection. (A) Variances of the mRNA level, the rate of translation, and the protein level across lineages through time. (B) End-point transcription-translation correlation across lineages. (C) End-point mRNA-protein correlation. Blue lines in (B) and (C) are least-squares regression lines. (D-E) End-point transcription-translation correlation (D) and mRNA-protein correlation (E) under different combinations of *N_e_* and SD of the fitness function. All phenotypes plotted are in log scale.

To confirm that a negative transcription-translation correlation arises due to stabilizing selection, we varied the strength of stabilizing selection by altering the effective population size *N_e_* and standard deviation of the fitness function *σ_ω_*. As expected, the transcription-translation correlation became more negative when selection was stronger (i.e., greater *N_e_* and/or smaller *σ_ω_*) (Fig. 1D). In contrast, the mRNA-protein correlation was close to zero when selection was strong, but more positive when selection was weak (Fig. 1E). Together, these observations support the view that compensatory evolution between the mRNA level and the translation rate helps maintain a relatively constant protein level when protein levels are subject to stabilizing selection.

We also conducted simulations along a phylogenetic tree of 50 species (Fig. 2A) and estimated evolutionary correlations between traits (i.e., the correlation accounting for phylogenetic history) to confirm whether the above patterns would be seen in phylogenetic comparative analyses. Consistent with observations from the simulated replicate lineages, the evolutionary correlation between the mRNA level and the translation rate is strongly negative when the protein level is under stabilizing selection (*r ≈ −*0.8 under the parameter combinations we considered, Table S2). As this correlation is not as negative as that across replicate lineages or across simulations (i.e., correlation calculated from same species’ phenotypes in 500 simulations, which is mathematically equivalent to 500 replicate lineages), we repeated the simulation along trees transformed using Pagel’s *λ* transformation [27] (i.e., extending external branches and shortening internal branches of the original tree) to test if the discrepancy is due to small effective sample size caused by the tree structure [28]. Indeed, evolutionary correlations estimated from the transformed trees were more similar to the correlations among replicate lineages (Table S3). We also observed a positive association of divergence time with both the mRNA level and the translation rate, while obvious saturation was seen for the protein level (Fig. 2B), consistent with the time-variance relationships seen in simulations along replicate lineages. Fitting standard phylogenetic models of continuous trait evolution (see Methods) indicates that divergence of the protein level is better described by an Ornstein–Uhlenbeck (OU) process, while divergence of the mRNA level and the translation rate are better described by a Brownian motion (BM) process (Fig. S2A, C, and E). However, when measurement error was unaccounted for, model comparisons to the mRNA level and the translation rate favored OU models as well (Fig. S2A, C, E). This is consistent with previous findings that measurement error biases model fits toward OU models even if the traits evolved under a BM model [29, 30].

**Figure 2:**
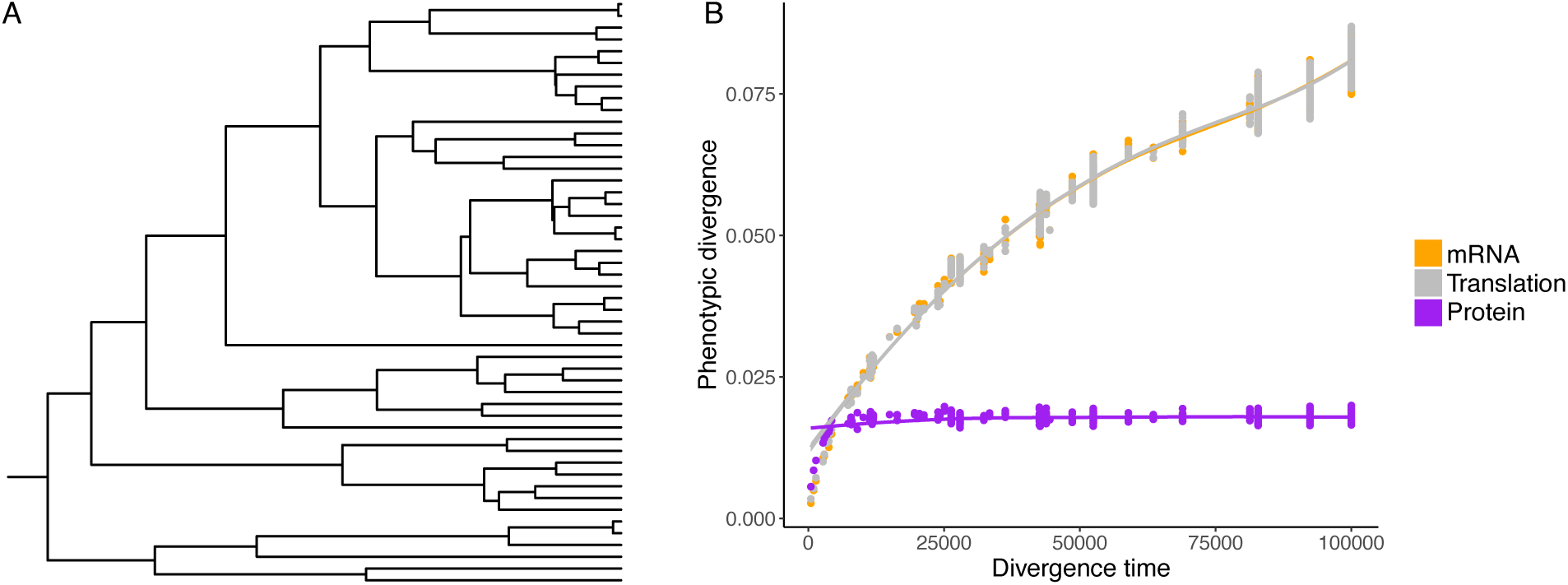
(A) Phylogenetic tree used for the simulation. The root edge is only shown to indicate the root’s location. (B) Pairwise phenotypic divergence plotted against pairwise divergence time when the protein level is under stabilizing selection. Each data point represents a combination of species pair and trait. The y-axis value of each point is the absolute phenotypic average difference between the two species (i.e., |ln *R_i_|*, |ln *β_i_|*, and |ln *P_i_|* for species *i* and *j*), averaged across 500 simulations. Each curve is a locally estimated scatterplot smoothing (LOESS) curve for the corresponding trait.

### Transcription-translation coevolution of interacting genes

Due to shared gene regulatory architecture, the expression levels of different genes are not independent of each other, and this interdependence can potentially shape the mutational architecture and influence the way evolution of expression levels is constrained [31]. Therefore, we also examined how interactions between genes influence the coevolution of mRNA and protein levels. We simulated data under a model of two interacting genes where each gene’s realized transcription rate is determined collectively by its genotypic value (i.e., a baseline transcription rate) and the regulatory effect of protein product(s) of other gene(s) (Fig. 3A, also see Methods).

**Figure 3:**
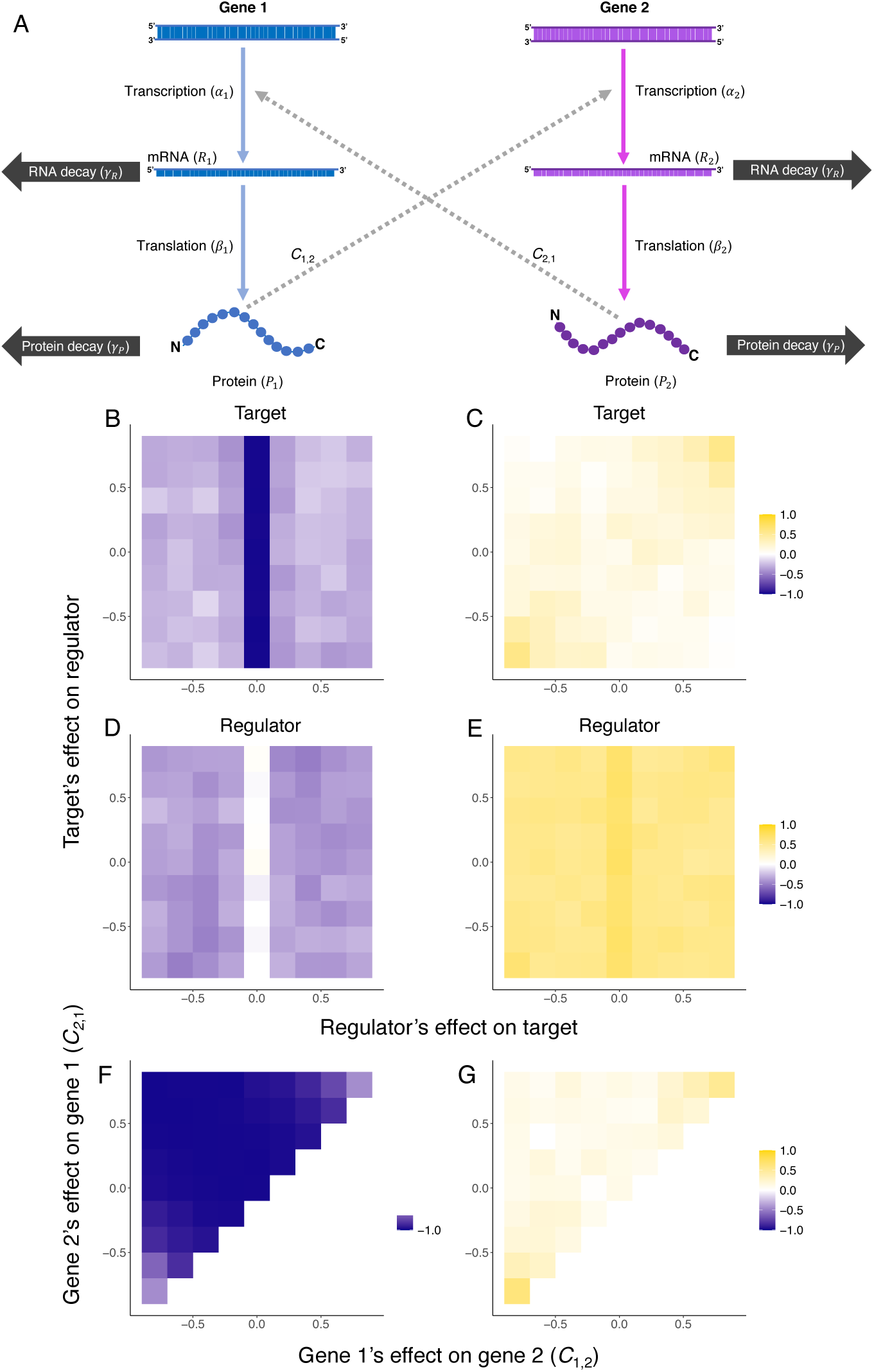
(A) Schematic illustration for the model of between-gene interaction considered in this study. (B-G) Transcription-translation correlation and mRNA-protein correlation of interacting genes. Axes are genes’ regulatory effects on each other. (B-C) A gene directly subject to stabilizing selection (i.e., has an optimal protein level). (D-E) A regulator gene that is not directly subject to selection. Transcription of the target gene in (B) and (C) is regulated by the protein of the regulator in (D) and (E). (F-G) Correlations observed for one gene (gene 1) when both genes are directly subject to stabilizing selection.

We first examined the scenario where one gene is only a regulator (referred to as “the regulator”) while the protein level of the other gene is subject to stabilizing selection (referred to as “the target”). The target gene, which is directly under stabilizing selection, showed negative transcription-translation correlation (i.e., correlation between genotypic values of transcription and translation; Fig. 3B) and weak mRNA-protein correlation (Fig. 3C) for most combinations of interaction parameter values. However, the negative transcription-translation correlation became weaker in the presence of the regulator, reflecting between-gene compensatory evolution. In other words, in the presence of a regulator, a deleterious substitution affecting the target’s transcription or translation rates can be compensated by a substitution affecting transcription or translation rate of either the target itself or the regulator. In the presence of strong negative feedback (i.e., regulatory effects of two genes on each other both have large absolute values but opposite signs), the mRNA-protein correlation became more positive. The regulator also exhibited a negative transcription-translation correlation (Fig. 3D), indicating that indirect selection due to the regulatory roles of a gene is sufficient to cause its transcription-translation compensatory evolution. The mRNA-protein correlation of the regulator was weakened by the interaction but remained rather strong across the parameter space (Fig. 3E). When both genes under consideration are directly subject to stabilizing selection, we observed a negative transcription-translation across the examined parameter space, yet was weaker in the presence of strong negative feedback (Fig. 3F). Consistently, the mRNA-protein correlation is generally weak but becomes stronger when there is strong negative feedback (Fig. 3G).

Although the number of genes involved in real regulatory networks is much greater, it is impractical to evenly sample a higher-dimensional space of interaction parameter combinations. To this end, we examined a series of motifs that bear features commonly seen in real regulatory networks (Fig. S4). The correlations were mostly consistent with those of genes in two-gene regulatory motifs: the regulator(s) and the target(s) all showed negative, though not necessarily strong transcription-translation correlations, while targets subject to direct stabilizing selection showed weaker mRNA-protein correlation (Table S4). Together, these results show transcription-translation compensatory evolution can take place as a result of a gene’s regulatory effect on other gene(s), yet would be less pronounced due to other gene(s)’ regulatory effect(s) on the gene of interest.

### Transcription-translation coevolution of functionally equivalent genes

Next, we considered a scenario where two genes express functionally equivalent proteins, such that fitness is determined by the total amount of proteins expressed from the two genes. These genes can be perceived as duplicate genes that have not yet been divergent enough in their protein sequences to be functionally distinct, in which case selection would act to maintain the total expression level [32]. In simulations under this scenario, both genes show negative transcription-translation correlations (Table S5, though the correlations are not as strong as that in the single gene case (Fig. 1A, S1). We also observed negative correlations between expression traits (i.e., mRNA levels, translation rates, and protein levels) of different genes (Table S5). This scenario, like examples of interacting genes (Fig. 3B, C, F, G), demonstrates that between-gene compensatory evolution can complement compensatory evolution of transcription and translation of the same gene and weaken the negative transcription-translation correlation.

### Transcription-translation coevolution under directional selection

To understand how directional selection might influence the coevolution of mRNA and protein levels differently, we simulated evolution towards the optimal protein level in 500 replicate lineages starting from the same phenotype (see Methods). The end-point protein level is distributed around the new optimum, yet the relative contribution of mRNA level and translation to evolutionary change in the protein level varied across lineages (Fig. 4A-B). The end-point mRNA level and the end-point translation rate are negatively correlated (Fig. 4A), while the protein level is essentially uncorrelated with the mRNA level (Fig. 4B). This was similar to the observed correlations when the protein level is under stabilizing selection.

**Figure 4:**
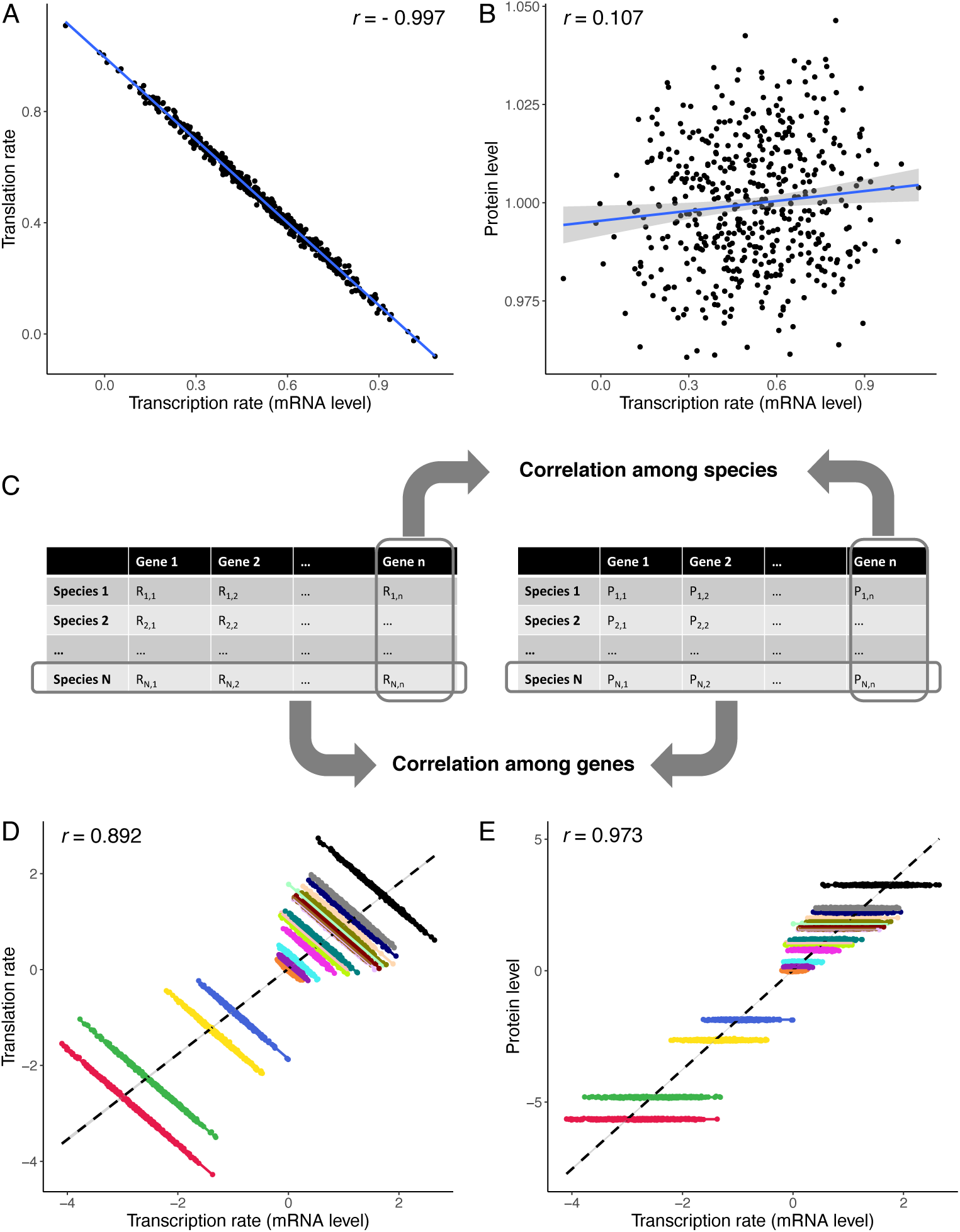
Coevolution of the mRNA level, the rate of translation, and the protein level when the protein level is under directional selection. (A) End-point transcription-translation correlation. (B) End-point mRNA-protein correlation. (C) A schematic illustration of the difference between across-species and across-gene correlations, using mRNA-protein correlation as an example. Correlation across species is calculated from mRNA and protein levels of the same gene in different species, whereas correlation across genes is calculated from mRNA and protein levels of different genes in the same species. (D) End-point transcription-translation correlation among multiple genes with different optimal protein levels. (E) End-point mRNA-protein correlation among multiple genes with different optimal protein levels. In (D) and (E), each gene is represented by a cloud of points of a distinct color. Solid lines of different colors are least-squares regression lines of different genes, while the dashed lines are least-squares regression lines based on all data points. Correlation coefficient shown in each panel is based on all data points in the panel.

As different genes within a genome likely have different optimal protein levels, we repeated the above simulations for multiple genes with the same starting mRNA levels and translation rates but different optimal protein levels. Specifically, we asked if patterns of correlation across species (within gene) and that across genes would be different (difference between two types of correlation illustrated in Fig. 4C). Patterns of correlations across species and across genes are drastically different: for each gene, there is a strong negative correlation between the mRNA level and the translation rate, but essentially no correlation between the mRNA level and the protein level. In contrast, both correlations are positive when compared among genes (Fig. 4D-E, Table S6). These observations reflect the fact that evolutionary changes in a gene’s mRNA level and translation rate were usually concordant (i.e., changing the protein level in the same direction), despite the negative correlation in terms of magnitude.

## Discussion

In this study, we demonstrate how the mRNA level, the translation rate, and the protein level coevolve when it is the protein level that is subject to selection. Using simulated data generated under a quantitative genetics model of gene expression evolution, we show that stabilizing selection on the protein level can cause a negative transcription-translation correlation (Fig. 1A) and weaken the mRNA-protein correlation (Fig. 1B). As measurement errors are known to impact empirical estimates of gene expression, we examined the impact of random error added to the end-point traits on our results. While measurement error weakened the negative transcription-translation correlation under neutrality, it does not account for the strong correlation observed in the presence of stabilizing selection on the protein level (Table S1). Notably, as errors in the translation rate are correlated with errors in the mRNA level, a spurious correlation between estimates of these two traits might arise. Our simulations also reveal that stabilizing selection on the protein level can make the protein level more conserved across species than the mRNA level (Fig. 1C, Fig. 2B), which is a pattern often found in empirical studies [10, 11, 17]. However, we note this phenomenon is only expected to occur under some combinations of evolutionary parameters (see the “Expected phenotypic variances” subsection of the Methods).

The data points in Fig. 1A, which correspond to different lineages, can also be interpreted as representing different genes. In this case, the end-point phenotypes shall be labeled as divergence from the ancestral phenotype. Such negative correlation between evolutionary divergence in transcription and translation of different genes has been observed in empirical studies of both budding yeasts [19, 20] and mammals [18]. The negative correlations observed in these empirical studies shown in were weaker than those observed in our simulations, likely because different genes have different evolutionary parameters (e.g., protein levels of different genes are not under equally strong selection).

We observed that the evolution of the mRNA level and the translation rate, which are not directly under selection and have no optima was qualitatively similar under both neutral evolution (i.e., no optimum for protein levels) and protein levels subject to stabilizing selection. Under neutrality, variance among lineages is expected to increase through time at a rate determined by the mutational variance [33, 34]. In our simulations where the protein level is under stabilizing selection, variances in the mRNA level and the translation rate among replicate lineages increased through time without saturation (Fig. 1C), and phylogenetic analysis of simulations along a phylogenetic tree favored a Brownian motion model as well, consistent with neutral expectations. Importantly, the evolution of the mRNA level and the translation rate were affected by selection, as they underwent much less evolutionary divergence than expected under neutrality. Such apparently (but not truly) neutral evolution occurs when a trait is not under direct selection but genetically correlated to trait(s) subjected to constraint: some mutations affecting the focal traits are purged by selection due to their deleterious effect on other trait(s), yet the focal trait itself has neither an optimal value nor boundaries, allowing it to diverge indefinitely, albeit slowly [35]. It should be noted that the mRNA level and the rate of translation are presumably not truly unbounded, as resources within the cell (i.e., RNA polymerase molecules, ribosomes, ATP, etc.) are ultimately limited, though we assumed that the phenotypes are far from such limits in our simulations.

Previous studies have shown evolutionary divergence in the mRNA level at phylogenetic scales is best described by an OU process [36–38], which appears in contradiction to our observation that variance in the mRNA level continued to increase through time (Fig. 1C, 2B). One possible explanation for this is that measurement errors create a bias in favor of the OU model [29, 30], which is confirmed in this study (Fig. S2). The discrepancy between our simulation results and observations from analyses of real data is likely to be due in part to measurement error. It is also worth noting that the same amount of error would cause more severe bias when the total amount of divergence is low. Therefore, stabilizing selection on the protein level does make it more likely that divergence in the mRNA level appears to fit an OU model by augmenting the influence of measurement errors. Note that there could be other, non-mutually exclusive explanations to these discrepancies. For example, there might be an upper bound to each gene’s mRNA level and translation rate imposed by the availability of cellular resources.

Directional selection on the protein level results in patterns of correlations that are similar to those resulting from stabilizing selection. To reach the optimal protein level, the mRNA level and the rate of translation undergo evolutionary changes that are concordant in terms of direction, but complementary in terms of magnitude. As a result, negative transcription-translation correlation and weak mRNA-protein correlation are both expected among a group of species that underwent selection towards the same optimal protein level. After the optimum is reached, stabilizing selection takes over and continues to promote the same kind of correlation. The effects of stabilizing and directional selection can be collectively viewed as the effect of the fitness landscape: when there exists an optimal protein level, selection is expected to result in a negative correlation between the mRNA level and the translation rate, and weak to no correlation between the mRNA level and the protein level.

The negative transcription-translation correlation and weak mRNA-protein correlation resulting from selection on the protein level are within gene and across species. We also extended our model to explore how the same evolutionary processes would shape the correlations across different genes. We show strongly positive transcription-translation and mRNA-protein correlations among genes with different optimal protein levels, demonstrating an instance of Simpson’s paradox (i.e., the correlation between variables seen in certain subsets of data differs from that seen in the complete dataset). Recent studies have found rather strong mRNA-protein correlations across genes (e.g., [7, 8]), and it is suggested that measurement errors played a significant role in weakening the correlations in earlier studies [26]. Our finding reconciles these results and the weaker with-gene, among-species correlations.

In our simulations, we considered a simple mutational architecture with no pleiotropic mutations (i.e., each mutation affects either transcription or translation but not both) as parameterization of pleiotropy in the simulations is challenging. The prevalence of pleiotropic mutations and their effects on transcription and translation are unclear. If the mRNA level and the rate of translation could be measured simultaneously for a sufficiently large number of mutant genotypes, a more complete picture of the two traits’ mutational architecture could be obtained, which would allow better parameterization. Similarly, we did not consider mutations affecting the degradation rates due to the difficulty of parameterization.

## Conclusion

By connecting patterns in the coevolution of the mRNA level, the rate of translation, and the protein level to explicit evolutionary processes, we demonstrate that several widely observed phenomena in between-species comparisons — namely, weak mRNA-protein correlation, negative transcription-translation correlation, and the protein levels being more evolutionarily conserved than the mRNA level — can all result from stabilizing selection on the protein level. Additionally, positive mRNA-protein correlations across genes arise because different genes have different optimal protein levels. With these connections built, our results can aid the interpretation of observation in future empirical studies and help disentangle the effects of biological and technical factors.

## Methods

### Model of gene expression

The basic model we study is a system of two equations. The first describes the rate at which the mRNA level of a single gene, *R* changes with time. This is given by

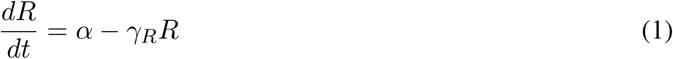

where *α* is the gene’s transcription rate, and *γ_R_* is the rate at which the mRNA molecules are degraded. The rate at which the protein level, *P* changes with time is described as

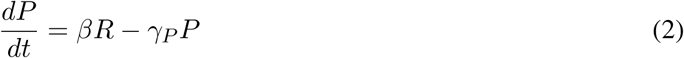

where *β* is the per-transcript translation rate, and *γ_P_* is the rate by which protein products are degraded *γ_P_*. The *β* parameter can be interpreted as the rate of translation initiation, as initiation has been shown to be a major rate-limiting step of translation [39]. At the equilibrium state (i.e., neither the mRNA level nor the protein level is changing),

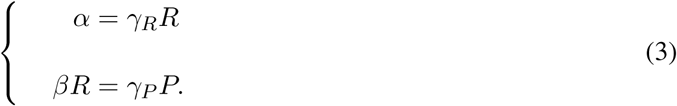

We can solve Eqn. (3) and obtain

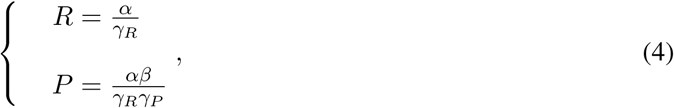

which can then be log-transformed

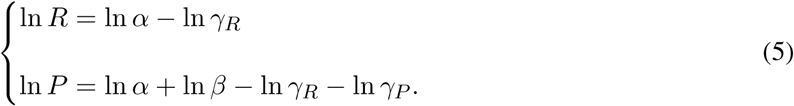

In our model, we assume a precise match between the genotypic values (i.e., *α* and *β*) and the phenotype (i.e., equilibrium *R* and *P*) and did not consider expression noise. Although expression noise could play a role in constraining evolution of transcription and translation rates [40], we assume, in this study, that selection imposed by noise is far weaker than that imposed by the mean protein level and thus negligible.

### Expected phenotypic variances

Under the assumption that degradation rates in Eqn. (5) are constant, the variance of the mRNA level across samples (i.e., replicate lineages) is equal to variance of the transcription rate (i.e., Var(ln *R*) = Var(ln *α*)). Variance of the protein level, as a function of other variances, is given by

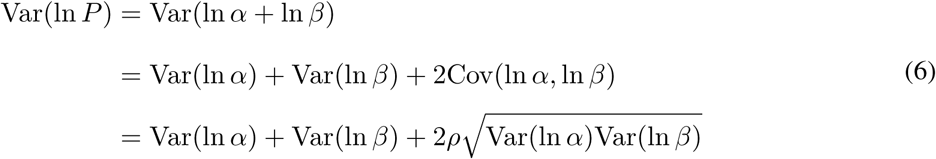

where Cov(ln *α,* ln *β*) is the covariance between ln *α* and ln *β*, and *ρ* is their correlation coefficient. The protein level is more conserved than the mRNA level if:

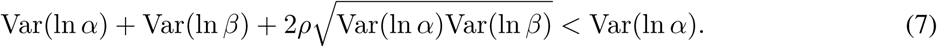

If we re-arrange the inequality, we obtain the conditions under which we would expect to see the more interlineage variation in *R* than *P*

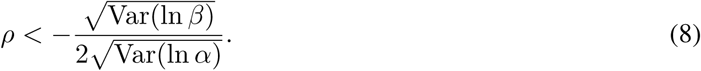

We note that the inequality in Eqn. (8) is invalid when 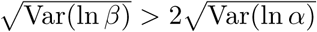 since *ρ* is only defined between -1 and 1, which means the protein level can be more conserved than the mRNA only if variance in the rate of translation is not too much higher than variance of the mRNA level. The variances (Var(ln *β*) and Var(ln *α*)) are depending on multiple factors, including strength of selection on the protein level, mutational parameters, and indirect effect of selection due to interaction with other gene(s) (see below). Therefore, the phenomenon the the protein level is more conserved than the mRNA level reflects the collective effect of multiple factors and may not occur depending on value of different parameters.

### Model of interaction between genes

Let us consider two interacting genes, gene 1 and gene 2. Gene 1’s protein can influence gene 2’s transcription, and vice versa. We assume that the expression level of gene 1 and gene 2 are governed by the dynamics depicted above, with each gene having its own genotypic values for the translation and transcription rate parameters (i.e., the *α* value for gene 1 is *α*_1_ and for gene 2 is *α*_2_). We assume the rates of mRNA and protein degradation are the same for all genes. In this model, the two genes interact according to a set of interaction parameters *C*. The parameter *C*_1,2_ is the effect of gene 1’s protein product *P*_1_ on gene 2’s transcription level, and *C*_2,1_ is the reverse. We can then set up a system of differential equations as follows:

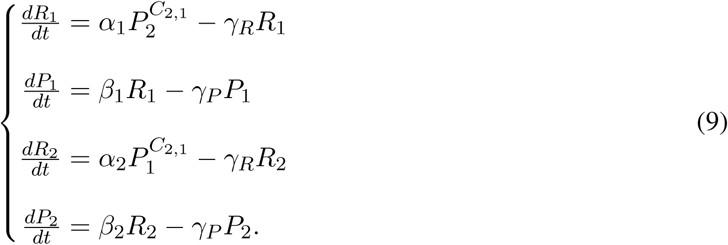

When the interaction parameter is negative, the regulatory effect on the target gene is repression. This is reflected in the asymptotic decrease of the realized transcription rate 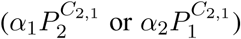 as the concentration of the repressor increases. Conversely, a positive interaction parameter indicates an activation effect, where the realized transcription rate increases with the concentration of the activator.

It is worth noting that the activation effect would plateau as the activator’s concentration increases, because an excess of activator molecules would not be able to bind the target. Although the realized transcription rate increases monotonously in our model, our approximation remains reasonable when the interaction parameter is between 0 and 1 and the regulator’s expression level is not extremely high. Given our focus on scenarios where stabilizing selection is in action, we can assume that the concentration of the regulator’s protein would never reach a level where target’s transcription rate plateaus. We did not consider interaction parameter values above one in this study.

At equilibrium, the following four equations hold:

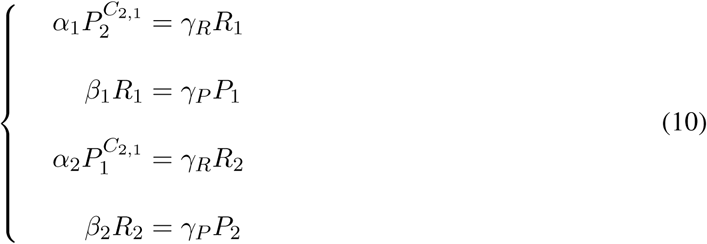

After log-transformation and rearrangement, we can express Eqn. (10) in matrix form as follows:

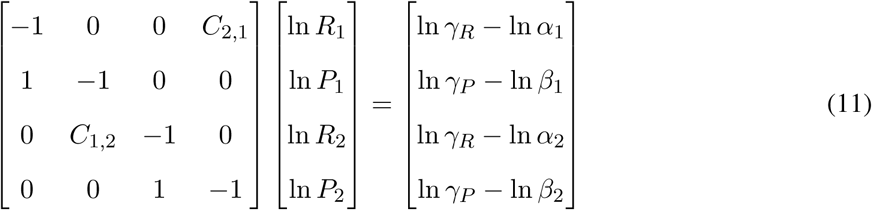

Solving the system of linear equations gives the logarithms of *R*_1_, *R*_2_, *P*_1_, and *P*_2_.

This model can be extended to systems of three or more genes. For gene *i* in a system of *n* genes, we have the following differential equations:

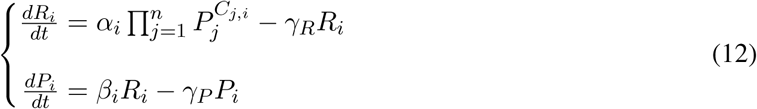

Again, after log-transformation and rearrangement, we can express Eqn. (12) in matrix form as follows:

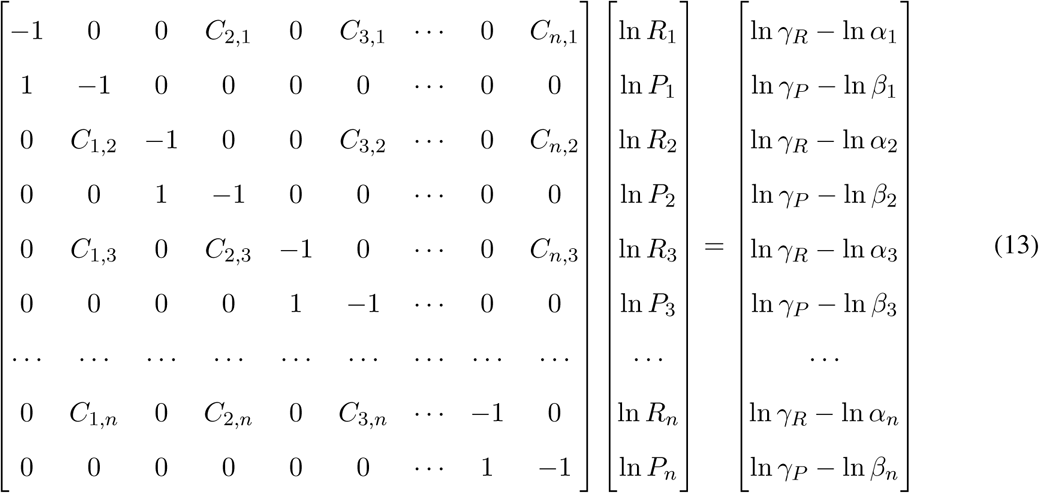

Note that the system may not have a solution (i.e., the leftmost matrix may not be invertible), depending on values of the parameters. A biological mechanism underlying such scenarios is positive feedback: if a set of genes activate each other, and their initial expression levels are high enough such that the degradation cannot counteract the increase of their expression, there would not be an equilibrium. Instead, their expression levels would increase indefinitely until a physical barrier is reached (e.g., limited by availability of ribosomes).

When we simulated evolution of two interacting genes (see below), we considered all combinations of the following values for *C*_1,2_ and *C*_2,1_: −0.8, −0.6, −0.4, −0.2, 0, 0.2, 0.4, 0.6, and 0.8. When we simulated evolution of three interacting genes, we had absolute values of all non-zero interaction parameters equal to 0.5. The set of triple-gene regulatory motif examined in this study were chosen because they bear features commonly seen in real regulatory networks [41, 42]. Specifically, motif 1 is a simple regulator chain, motif 2 is a negative feedback loop, motifs 3-5 represent regulatory motifs where the same target is regulated by more than one regulators, while motifs 6-8 represent motifs where one regulator (i.e., transcription factor) regulates more than one targets (see Fig. S4 for graphical depictions of the motifs).

### Simulation of evolution along a single lineage

For a system of *n* genes, we considered a total of 2*n* traits that are directly affected by mutations, including log-transformed genotypic values of their transcription rates (ln *α*_1_, …, ln *α_n_*) and translation rates (ln *β*_1_, …, ln *β_n_*), and simulated the evolution of the population mean phenotype through time. Traits that are directly under selection are the equilibrium protein levels (ln *P*_1_, …, ln *P_n_*). Given the genotypic values, the steady-state protein levels are calculated using equation system (5) when only a single gene is under concern, using (11) for a system of two genes, or (13) for a system of three or more genes. As the degradation terms in Eqn. (5) are constants, we omitted them in the simulations when only a single gene is considered; that is, ln *R* is represented by ln *α*, and ln *P* is represented by ln *α* + ln *β*. The total number of mutations that would occur in time step *t*, denoted *m_t_*, is drawn from a Poisson distribution. The mean of the distribution, *E*[*m_t_*], is given by

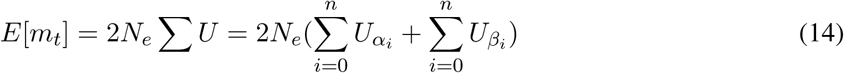

where *N_e_* is the effective population size, ∑*U* is the total per-genome rate of mutations affecting the transcription and translation rates, while *U_αi_* and *U_βi_* are rates of mutations affecting gene *i* ’s transcription and translation rates (i.e., number of mutations per haploid genome per time step), respectively. It is assumed here that a mutation can affect either transcription or translation rate of one gene, but not both.

Each mutation can affect either transcription or translation rate. The probability that a mutation affects the transcription rate of gene *i* is *U_αi_ /*∑*U*, and the probability that it affects the translation rate of gene *i* is *U_βi_ /*∑*U*. The phenotypic effect of a mutation affecting a gene *i*’s transcription rate (ln *α_i_*) is drawn from a normal distribution *𝒩* (0*, σ_αi_*). Similarly, if the mutation affects the translation rate of gene *i* (ln *β_i_*), its effect is drawn from another normal distribution, *𝒩* (0, *σ_βi_*). Given the protein level of a gene *i*, the fitness *ω* with respect to its protein level is given by a Gaussian function:

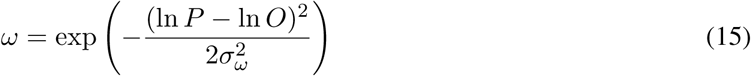

where *O* is the optimal protein level and *σ_ω_* is the standard deviation of the fitness function (also referred to as width of the fitness function).

In a system where protein levels of *n* genes are subject to selection, the overall fitness is calculated as:

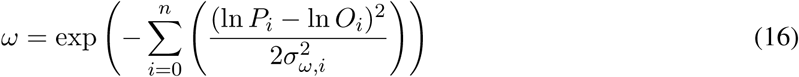

When *n* = 1, equation (16) gives the same result as (15). If there is any gene that has no equilibrium phenotype, the fitness would be treated as zero, though such situations did not occur in our simulations. The coefficient of selection, *s*, is then calculated as *s* = (*ω/ω_A_*) − 1, where *ω_A_*is the ancestral fitness. The fixation probability *p_f_* of the mutation is calculated following the approach of Kimura [43]:

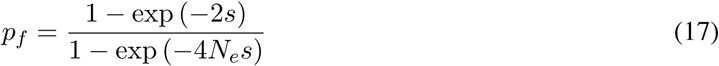

With probability *p_f_*, the mutation’s phenotypic effect would be added to the population’s mean before the next mutation is considered.

The above process would be repeated for *T* times for each lineage. We set *T* = 10^5^, *N_e_* = 1000, *U_α_* = *U_β_* = 5 *×* 10*^−^*^4^ (i.e., 2*N_e_U* = 1), *σ_α_* = *σ_β_* = 0.1, ln *O* = 0, and *σ_ω_* = 1 for all simulations, unless specified. When we simulated a single gene that is subject to directional selection, we set ln *O* = 1. All simulations started with ln *α* = 0 and ln *β* = 0, unless specified. For each combination of parameter values, we simulated 500 independent lineages.

For simulations where multiple genes with different optimal protein levels were involved, we randomly sampled a set of 20 optima ln *O ∼ 𝒩* (0, 2). For each gene, we conducted simulation along 500 replicate lineages. We assumed no linkage between these genes and had them evolve independently (i.e., simulations for different genes were run separately).

For simulations of functional equivalent genes, fitness is calculated as

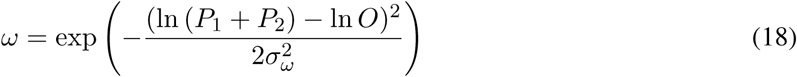

where *P*_1_ and *P*_2_ are two genes’ respective protein levels. In these simulations, the two genes are assumed to be unlinked and independently regulated (i.e., each mutation can affect either transcription or translation of only one gene).

It should be noted that our simulations were based on a sequential-fixation model of evolution, which allows more efficient simulations. That is, only one mutation is considered each time, and fixation probability of a mutation is calculated after it is determined if the previous mutation being considered is fixed. Such a model can be a good approximation as long as the mutation rate is not too high such that the probability that multiple mutations affecting the trait of interest segregate in the population at the same time is very low [44].

### Simulation along a phylogenetic tree

We generated a Yule tree (no extinction) and used this for all subsequent simulations (Fig. 2A). We then rescaled the tree to make its height (i.e., distance between the root and each tip) equal to 10^5^. We simulated evolution along each branch following the same procedure as above. The number of time steps is equal to the branch length (rounded down to the nearest integer). As this is purely for illustrative purposes (the model is the same as the case studied above), we used a single, representative tree. The qualitative patterns did not depend on the shape of the tree.

For each simulation, we estimated the evolutionary variance-covariance (VCV) matrix between lineages. To assess the influence of tree structure, we used a *λ* transformation [27] to rescale the relative length of terminal branches with tree height kept the same. We fitted two models, Brownian motion (BM) and Ornstein-Uhlenbeck (OU) to each trait (the mRNA level, the translation rate, and the protein level) for each simulation and compared the relative support of the models using their sample-size corrected AICc weights (following [29]) and averaged the AICc weights across replicate simulations. All phylogenetic analyses were conducted using geiger [45].

### Adding measurement errors to simulated data

For each sample (i.e., an independent lineage or a tip of the phylogenetic tree), we added errors to the end-point mRNA and protein levels; the errors *ε* were normally distributed and centered on the true value (i.e., *ε ∼ 𝒩* (0*, σ_ɛ_*)). We considered values of *σ_ɛ_* = 0.01, 0.02, 0.03, 0.04, 0.05 in this study for both mRNA and protein levels. The “estimated” translation rate is calculated as a ratio of the “measured” protein and mRNA levels, such that error in the translation rate is correlated with error in the mRNA level.

### Code availability

All R code is available at https://github.com/phylo-lab-usc/Expression_Evolution.

## Acknowledgements

We thank members of the Pennell, Edge, and Mooney labs for their thoughtful comments on this study. M.P. and D.J. were supported by startup funds from the University of Southern California. J.Z. was funded by the grant R35GM139484 from the NIH. A.L.C. was funded by a NIH-IRACDA postdoctoral fellowship from Rutgers University. ChatGPT helped improve the clarity of the text.

## Supplementary materials

**Table S1:** Transcription-translation correlation and mRNA-protein correlation across replicate lineages under different levels of measurement error.

| Error SD | Selection regime | Correlation coefficient |  |
| --- | --- | --- | --- |
|  |  | Transcription-translation | mRNA-protein |
| 0 | Stabilizing selection | -0.9765 | 0.09209 |
|  | Neutral | 0.0004967 | 0.7126 |
| 0.01 | Stabilizing selection | -0.9670 | 0.05904 |
|  | Neutral | 0.0003159 | 0.7121 |
| 0.02 | Stabilizing selection | -0.9487 | 0.06802 |
|  | Neutral | 0.003284 | 0.7139 |
| 0.03 | Stabilizing selection | -0.9216 | 0.05275 |
|  | Neutral | -0.002881 | 0.7107 |
| 0.04 | Stabilizing selection | -0.8714 | 0.1294 |
|  | Neutral | -0.007650 | 0.7072 |
| 0.05 | Stabilizing selection | -0.8570 | 0.02024 |
|  | Neutral | -0.02017 | 0.7057 |

**Table S2:**
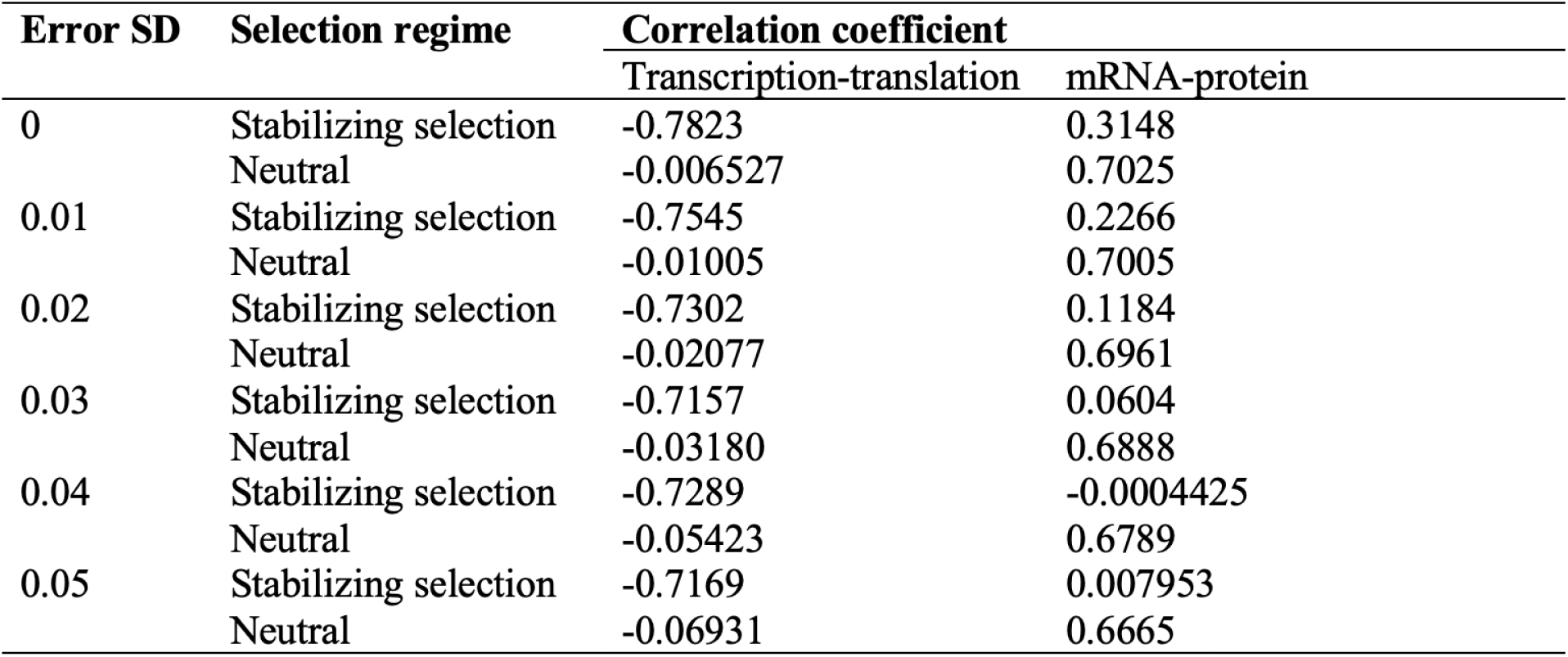
Evolutionary correlations between traits under different levels of measurement error.

| Error SD | Selection regime | Correlation coefficient |  |
| --- | --- | --- | --- |
|  |  | Transcription-translation | mRNA-protein |
| 0 | Stabilizing selection | -0.7823 | 0.3148 |
|  | Neutral | -0.006527 | 0.7025 |
| 0.01 | Stabilizing selection | -0.7545 | 0.2266 |
|  | Neutral | -0.01005 | 0.7005 |
| 0.02 | Stabilizing selection | -0.7302 | 0.1184 |
|  | Neutral | -0.02077 | 0.6961 |
| 0.03 | Stabilizing selection | -0.7157 | 0.0604 |
|  | Neutral | -0.03180 | 0.6888 |
| 0.04 | Stabilizing selection | -0.7289 | -0.0004425 |
|  | Neutral | -0.05423 | 0.6789 |
| 0.05 | Stabilizing selection | -0.7169 | 0.007953 |
|  | Neutral | -0.06931 | 0.6665 |

**Table S3:**
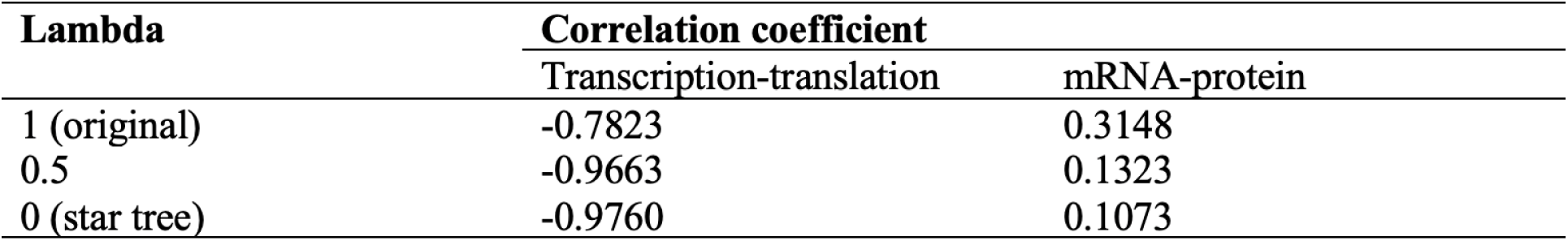
Evolutionary correlations estimated from results of simulations along transformed trees.

| **Lambda** | **Correlation coefficient** |  |
| --- | --- | --- |
|  | Transcription-translation | mRNA-protein |
| 1 (original) | -0.7823 | 0.3148 |
| 0.5 | -0.9663 | 0.1323 |
| 0 (star tree) | -0.9760 | 0.1073 |

**Table S4:** Transcription-translation correlation and mRNA-protein correlation of genes in triple-gene regulatory motifs. Red numbers indicate the gene’s protein level is directly subject to stabilizing selection.

| Regulatory motif | Correlation coefficient |  |  |  |  |  |
| --- | --- | --- | --- | --- | --- | --- |
|  | Transcription-translation |  |  | mRNA-protein |  |  |
|  | Gene 1 | Gene 2 | Gene 3 | Gene 1 | Gene 2 | Gene 3 |
| Motif 1 | -0.134 | -0.261 | -0.0848 | 0.651 | 0.670 | 0.134 |
| Motif 2 | -0.109 | -0.204 | -0.274 | 0.657 | 0.636 | 0.189 |
| Motif 3 | -0.236 | -0.271 | -0.249 | 0.609 | 0.600 | 0.101 |
| Motif 4 | -0.325 | -0.313 | -0.169 | 0.580 | 0.582 | 0.134 |
| Motif 5 | -0.266 | -0.208 | -0.108 | 0.886 | 0.897 | 0.135 |
| Motif 6 | -0.406 | -0.493 | -0.511 | 0.579 | 0.107 | 0.139 |
| Motif 7 | -0.536 | -0.536 | -0.485 | 0.491 | 0.0356 | 0.109 |
| Motif 8 | -0.368 | -0.330 | -0.368 | 0.312 | 0.286 | 0.279 |

**Table S5:**
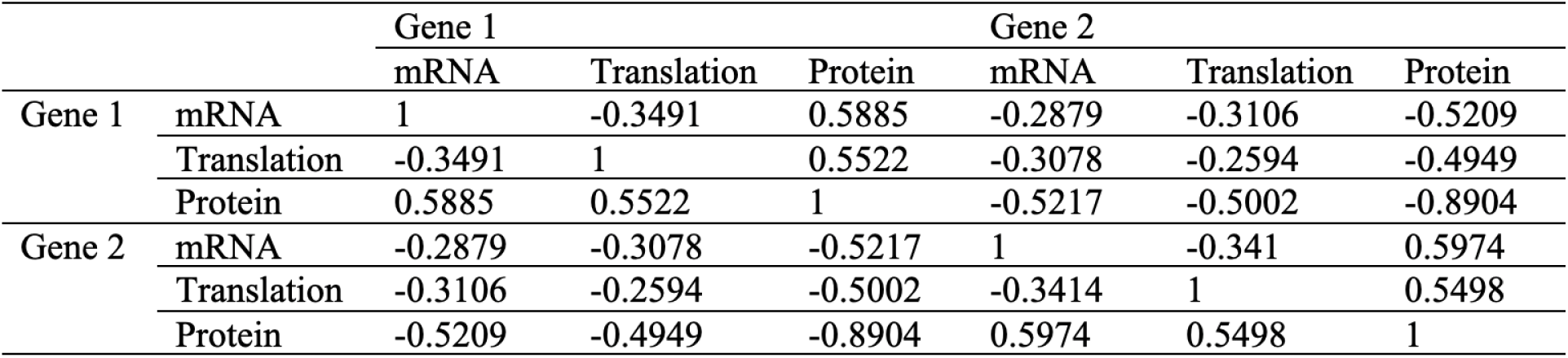
Correlation matrix for mRNA levels, translation rates and protein levels of two functionally equivalent genes (i.e., fitness is determined by the sum of two genes’ protein levels).

|  |  | Gene 1 |  |  | Gene 2 |  |  |
| --- | --- | --- | --- | --- | --- | --- | --- |
|  |  | mRNA | Translation | Protein | mRNA | Translation | Protein |
| Gene 1 | mRNA | 1 | -0.3491 | 0.5885 | -0.2879 | -0.3106 | -0.5209 |
|  | Translation | -0.3491 | 1 | 0.5522 | -0.3078 | -0.2594 | -0.4949 |
|  | Protein | 0.5885 | 0.5522 | 1 | -0.5217 | -0.5002 | -0.8904 |
| Gene 2 | mRNA | -0.2879 | -0.3078 | -0.5217 | 1 | -0.341 | 0.5974 |
|  | Translation | -0.3106 | -0.2594 | -0.5002 | -0.3414 | 1 | 0.5498 |
|  | Protein | -0.5209 | -0.4949 | -0.8904 | 0.5974 | 0.5498 | 1 |

**Table S6:** Transcription-translation correlation and mRNA-protein correlation of genes with different optimal protein levels.

| Gene # | Optimal protein level<br>(ln *O*) | Correlation coefficient |  |
| --- | --- | --- | --- |
|  |  | Transcription-translation | mRNA-protein |
| 1 | -5.653 | -0.9995 | 0.06265 |
| 2 | -4.814 | -0.9994 | 0.1049 |
| 3 | -2.643 | -0.9986 | 0.003202 |
| 4 | -1.869 | -0.9981 | 0.06626 |
| 5 | 0.006126 | -0.9734 | 0.1348 |
| 6 | 0.1369 | -0.9866 | 0.08257 |
| 7 | 0.3293 | -0.9930 | 0.09362 |
| 8 | 0.7829 | -0.9958 | -0.07563 |
| 9 | 0.9896 | -0.9971 | 0.06707 |
| 10 | 1.050 | -0.9973 | 0.09356 |
| 11 | 1.177 | -0.9971 | 0.08134 |
| 12 | 1.581 | -0.9980 | 0.06722 |
| 13 | 1.611 | -0.9983 | 0.01392 |
| 14 | 1.680 | -0.9983 | 0.02384 |
| 15 | 1.788 | -0.9982 | 0.07870 |
| 16 | 1.862 | -0.9983 | -0.003530 |
| 17 | 2.009 | -0.9983 | 0.01340 |
| 18 | 2.218 | -0.9984 | 0.007559 |
| 19 | 2.379 | -0.9986 | 0.02051 |
| 20 | 3.258 | -0.9989 | -0.006015 |

**Figure S1:**
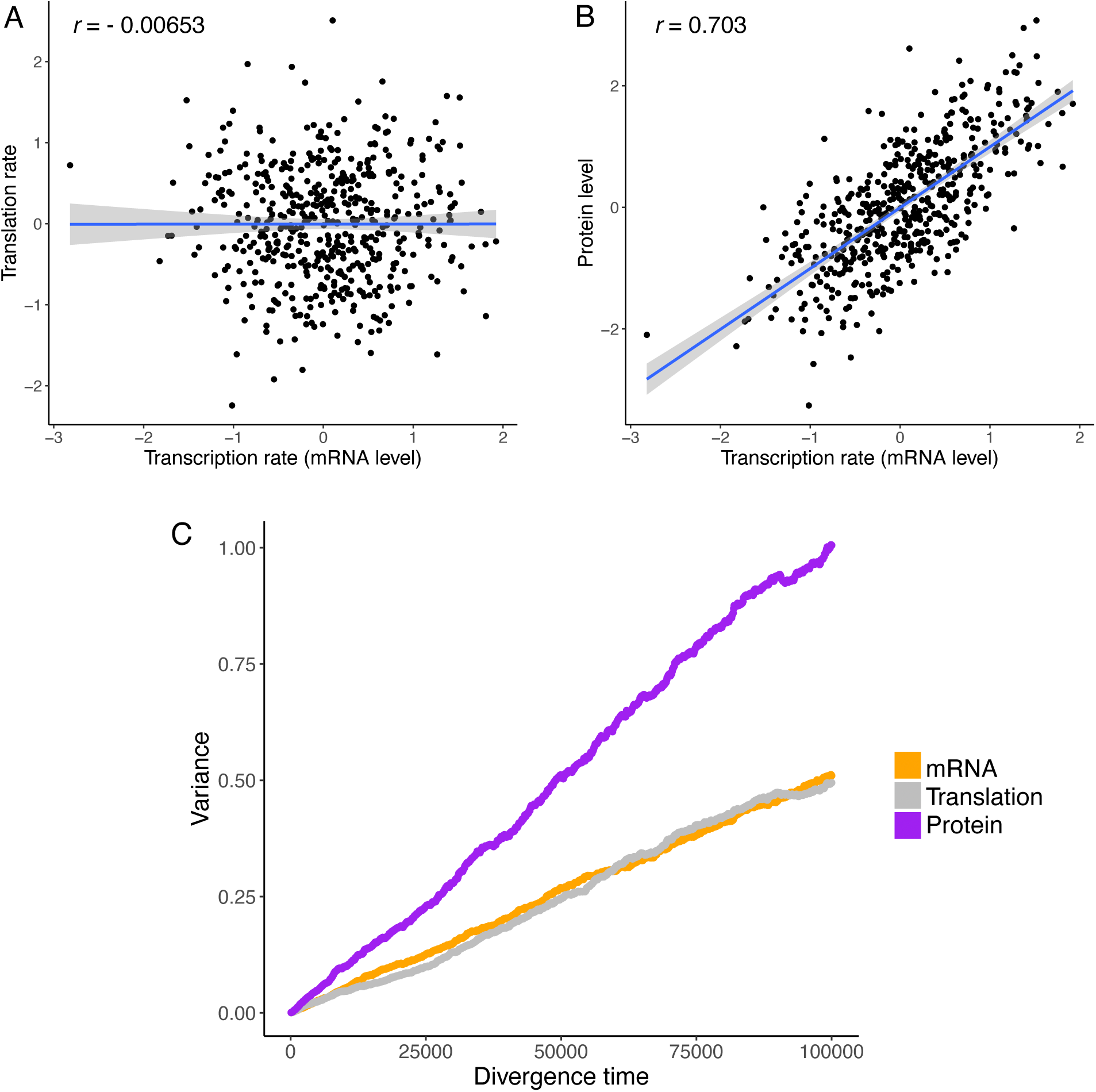
Coevolution of the mRNA level, the rate of translation, and the protein level under neutrality. (A) Variances of the mRNA level, the translation rate, and the protein level through time. (B) End-point correlation between the mRNA level and the translation rate. (C) End-point correlation between the mRNA level and the translation level. Blues lines in (B) and (C) are least-squares regression lines.

**Figure S2:**
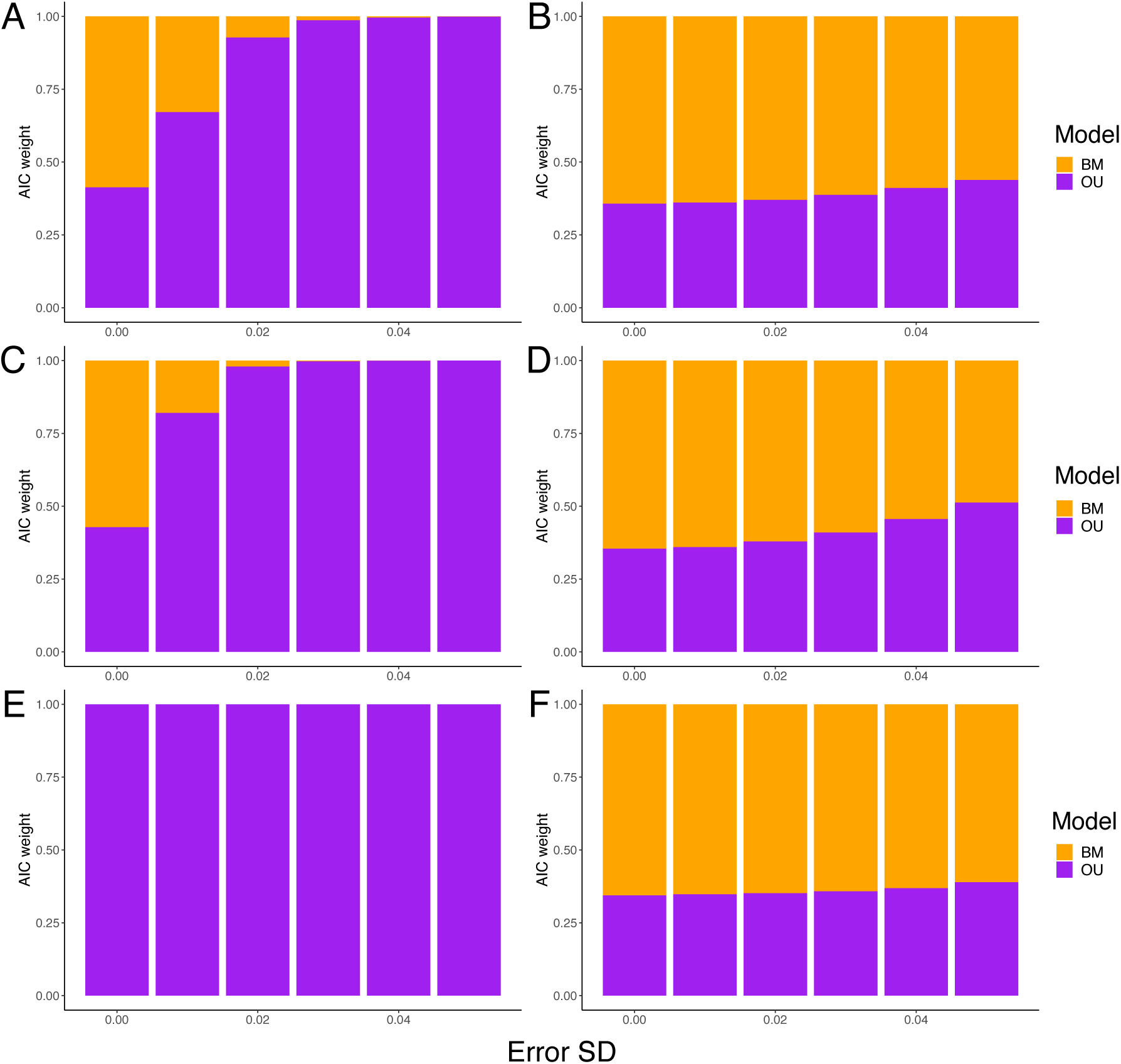
Relative support for Brownian motion (BM) and Ornstein-Uhlenbeck (OU) models when results of simulations along the phylogenetic tree in Fig. 2A, with tip phenotypes subject to different levels of measurement error. For each setting, the extent to which each model is supported is represented by the average AIC weight across 500 independent simulations. (A), (C), and (E) are for results of simulations where the protein level is under stabilizing selection, while (B), (D), and (F) are for simulations of neutral evolution. (A-B) AIC weights computed from the mRNA levels. (C-D) AIC weights computed from the translation rates. (E-F) AIC weights computed from the protein levels.

**Figure S3:**
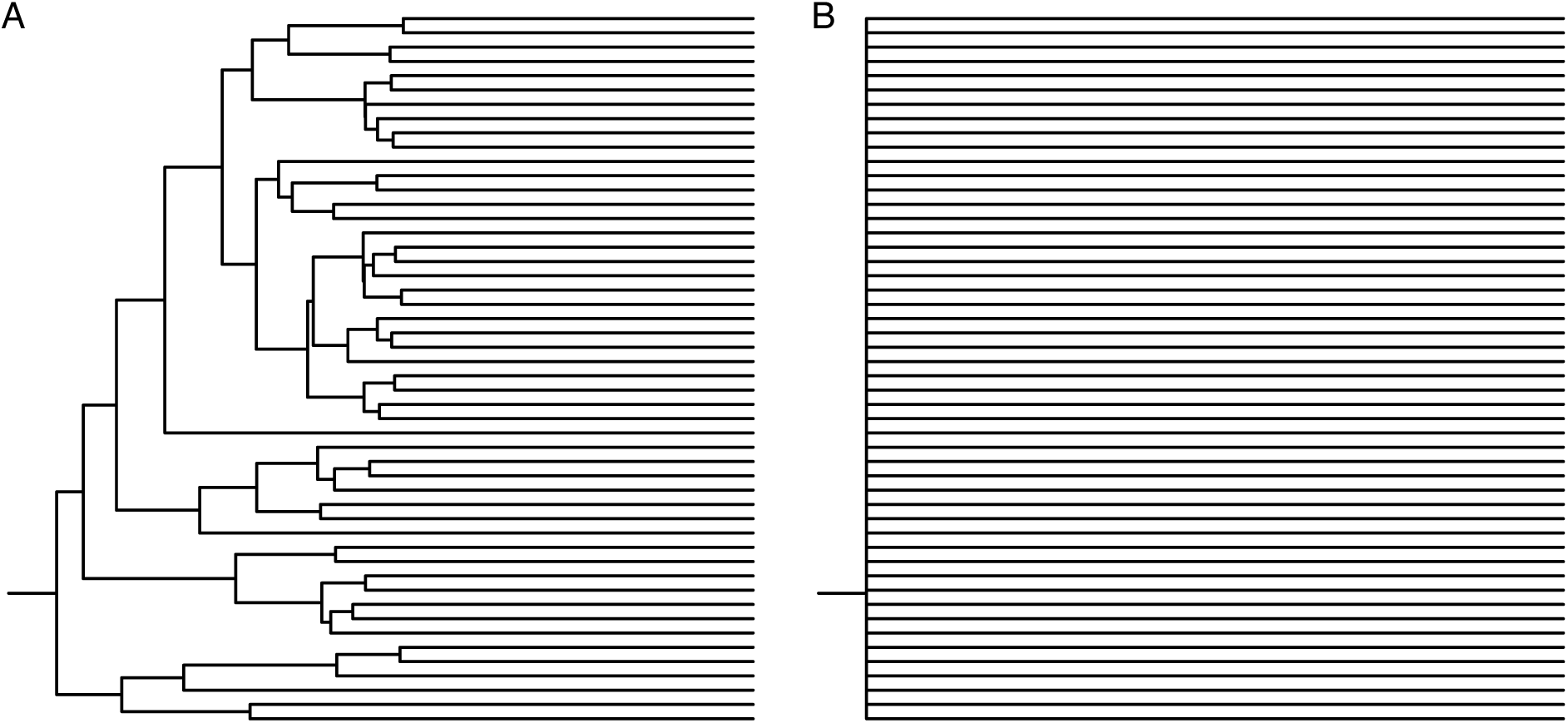
*λ*-transformed trees used in this study. (A) Tree transformed with *λ* = 0.5. (B) Tree transformed with *λ* = 0. The root edge is only shown to indicate the root’s location. The original tree is shown in Fig. 2A.

**Figure S4:**
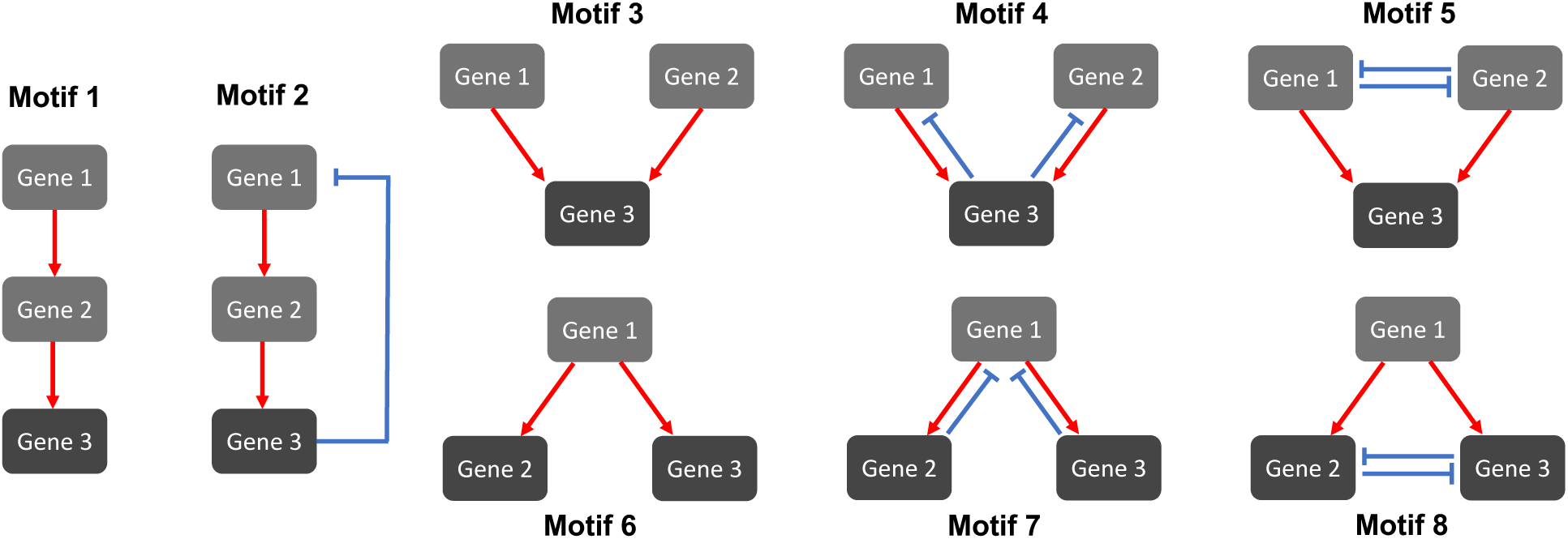
Special cases of triple-gene regulatory network motifs considered in this study. Genes whose protein levels are under direct selection (gene 3 in motifs 1-5, gene 2 and 3 in motifs 6-8) are colored dark gray, while regulators not subject to direct selection are colored light gray. Red arrows represent activation effects (*C_i,j_* = 0.5), while blue, flat-headed arrows represent repression effects (*C_i,j_* = −0.5).

## References

[1] King, M.-C & Wilson, A. C. (1975) Science 188, 107–116.

[2] Gygi, S. P, Rochon, Y, Franza, B. R, & Aebersold, R. (1999) Molecular and cellular biology 19, 1720–1730.

[3] Marguerat, S, Schmidt, A, Codlin, S, Chen, W, Aebersold, R, & Bähler, J. (2012) Cell 151, 671–683.

[4] Becker, K, Bluhm, A, Casas-Vila, N, Dinges, N, Dejung, M, Sayols, S, Kreutz, C, Roignant, J.-Y, Butter, F, & Legewie, S. (2018) Nature communications 9, 4970.

[5] Wang, D, Eraslan, B, Wieland, T, Hallström, B, Hopf, T, Zolg, D. P, Zecha, J, Asplund, A, Li, L.-h, Meng, C, et al. (2019) Molecular systems biology 15, e8503.

[6] Edfors, F, Danielsson, F, Hallström, B. M, Käll, L, Lundberg, E, Pontén, F, Forsström, B, & Uhlén, M. (2016) Molecular systems biology 12, 883.

[7] Franks, A, Airoldi, E, & Slavov, N. (2017) PLoS computational biology 13, e1005535.

[8] Fortelny, N, Overall, C. M, Pavlidis, P, & Freue, G. V. C. (2017) Nature 547, E19–E20.

[9] Perl, K, Ushakov, K, Pozniak, Y, Yizhar-Barnea, O, Bhonker, Y, Shivatzki, S, Geiger, T, Avraham, K. B, & Shamir, R. (2017) BMC genomics 18, 1–14.

[10] Laurent, J. M, Vogel, C, Kwon, T, Craig, S. A, Boutz, D. R, Huse, H. K, Nozue, K, Walia, H, Whiteley, M, Ronald, P. C, et al. (2010) Proteomics 10, 4209–4212.

[11] Khan, Z, Ford, M. J, Cusanovich, D. A, Mitrano, A, Pritchard, J. K, & Gilad, Y. (2013) Science 342, 1100–1104.

[12] Ba, Q, Hei, Y, Dighe, A, Li, W, Maziarz, J, Pak, I, Wang, S, Wagner, G. P, & Liu, Y. (2022) Science Advances 8, eabn0756.

[13] de Sousa Abreu, R, Penalva, L. O, Marcotte, E. M, & Vogel, C. (2009) Molecular BioSystems 5, 1512–1526.

[14] Vogel, C & Marcotte, E. M. (2012) Nature reviews genetics 13, 227–232.

[15] Liu, Y, Beyer, A, & Aebersold, R. (2016) Cell 165, 535–550.

[16] Buccitelli, C & Selbach, M. (2020) Nature Reviews Genetics 21, 630–644.

[17] Schrimpf, S. P, Weiss, M, Reiter, L, Ahrens, C. H, Jovanovic, M, Malmström, J, Brunner, E, Mohanty, S, Lercher, M. J, Hunziker, P. E, et al. (2009) PLoS biology 7, e1000048.

[18] Wang, Z.-Y, Leushkin, E, Liechti, A, Ovchinnikova, S, Mößinger, K, Brüning, T, Rummel, C, Grützner, F, Cardoso-Moreira, M, Janich, P, et al. (2020) Nature 588, 642–647.

[19] Artieri, C. G & Fraser, H. B. (2014) Genome research 24, 411–421.

[20] McManus, C. J, May, G. E, Spealman, P, & Shteyman, A. (2014) Genome research 24, 422–430.

[21] Long, S. B. (1980) Sociological methodology 11, 37–67.

[22] Fraser, H. B. (2019) Trends in Genetics 35, 3–5.

[23] Ho, W.-C & Zhang, J. (2019) Molecular biology and evolution 36, 604–612.

[24] Albert, F. W, Muzzey, D, Weissman, J. S, & Kruglyak, L. (2014) PLoS genetics 10, e1004692.

[25] Wang, S. H, Hsiao, C. J, Khan, Z, & Pritchard, J. K. (2018) Genome biology 19, 1–15.

[26] Csárdi, G, Franks, A, Choi, D. S, Airoldi, E. M, & Drummond, D. A. (2015) PLoS genetics 11, e1005206.

[27] Pagel, M. (1999) Nature 401, 877–884.

[28] Ané, C. (2008) The Annals of Applied Statistics 2, 1078–1102.

[29] Pennell, M. W, FitzJohn, R. G, Cornwell, W. K, & Harmon, L. J. (2015) The American Naturalist 186, E33–E50.

[30] Cooper, N, Thomas, G. H, Venditti, C, Meade, A, & Freckleton, R. P. (2016) Biological Journal of the Linnean Society 118, 64–77.

[31] Hether, T. D & Hohenlohe, P. A. (2014) Evolution 68, 950–964.

[32] Qian, W, Liao, B.-Y, Chang, A. Y.-F, & Zhang, J. (2010) Trends in Genetics 26, 425–430.

[33] Lynch, M & Hill, W. G. (1986) Evolution 40, 915–935.

[34] Khaitovich, P, Paabo, S, & Weiss, G. (2005) Genetics 170, 929–939.

[35] Jiang, D & Zhang, J. (2020) Evolution 74, 2158–2167.

[36] Bedford, T & Hartl, D. L. (2009) Proceedings of the National Academy of Sciences 106, 1133–1138.

[37] Chen, J, Swofford, R, Johnson, J, Cummings, B. B, Rogel, N, Lindblad-Toh, K, Haerty, W, Di Palma, F, & Regev, A. (2019) Genome research 29, 53–63.

[38] Dimayacyac, J. R, Wu, S, & Pennell, M. (2023) bioRxiv p. 2023.02.09.527893.

[39] Shah, P, Ding, Y, Niemczyk, M, Kudla, G, & Plotkin, J. B. (2013) Cell 153, 1589–1601.

[40] Hausser, J, Mayo, A, Keren, L, & Alon, U. (2019) Nature communications 10, 68.

[41] Lee, T. I, Rinaldi, N. J, Robert, F, Odom, D. T, Bar-Joseph, Z, Gerber, G. K, Hannett, N. M, Harbison, C. T, Thompson, C. M, Simon, I, et al. (2002) science 298, 799–804.

[42] Shen-Orr, S. S, Milo, R, Mangan, S, & Alon, U. (2002) Nature genetics 31, 64–68.

[43] Kimura, M. (1962) Genetics 47, 713.

[44] Lynch, M. (2020) Proceedings of the National Academy of Sciences 117, 10435–10444.

[45] Pennell, M. W, Eastman, J. M, Slater, G. J, Brown, J. W, Uyeda, J. C, FitzJohn, R. G, Alfaro, M. E, & Harmon, L. J. (2014) Bioinformatics 30, 2216–2218.

